# Microbiome derived acidity protects against microbial invasion in *Drosophila*

**DOI:** 10.1101/2023.01.12.523836

**Authors:** Alexander J. Barron, Danielle N. A. Lesperance, Jeremy Doucette, Sthefany Calle, Nichole A. Broderick

## Abstract

Microbial invasions underlie host-microbe interactions that result in microbial pathogenesis and probiotic colonization. While these processes are of broad interest, there are still gaps in our understanding of the barriers to entry and how some microbes overcome them. In this study, we explore the effects of the microbiome on invasions of foreign microbes in *Drosophila melanogaster*. We demonstrate that gut microbes *Lactiplantibacillus plantarum* and *Acetobacter tropicalis* improve survival during invasion of a lethal gut pathogen and lead to a reduction in microbial burden. Using a novel multi-organism interactions assay, we report that *L. plantarum* inhibits the growth of three invasive Gram-negative bacteria, while *A. tropicalis* prevents this inhibition. A series of *in vitro* and *in vivo* experiments revealed that inhibition by *L. plantarum* is linked to its ability to acidify both internal and external environments, including culture media, fly food, and the gut itself, while *A. tropicalis* diminishes the inhibition by quenching acids. We propose that acid produced by the microbiome serves as an important gatekeeper to microbial invasions, as only microbes capable of tolerating acidic environments can colonize the host. The methods described herein will add to the growing breadth of tools to study microbe-microbe interactions in broad contexts.

## Introduction

Animals consistently interact with complex communities of microorganisms throughout their lives. The collection of microbes that inhabits a particular niche is known as the microbiome. While many foreign microbes encountered by a host are transient, others have the potential to enter existing communities and establish themselves on or within the host, in a process known as microbial invasion^1,2^. Invasion can affect the host in different ways, depending on the actions of the microbe and the response by the host^3^. Positive outcomes of microbial invasion are observed during therapeutic fecal microbiome transplants, in which the microbiome from a healthy individual is transferred to a patient with gut disease, leading to a remodeling of the microbial community and improved health^4^. Conversely, microbial invasion can also produce negative outcomes on a host, as is the case during intestinal pathogenesis of *Escherichia coli* O157:H7^5^. Though the host outcomes are different in these cases, there is overlap in that new microbes enter and establish themselves within an existing community structure. The study of microbial invasion as a generalized concept is a relatively recent appreciation. The ability of a microbe to invade a community is generally governed by the introduction, establishment, and growth of foreign microbes in a new niche^1^. Attaining each phase of invasion requires that a microorganism overcome multiple barriers to entry, including physical and chemical barriers such as temperature and pH as well as biological barriers such as competition with the existing microbial community and actions of the host immune response^1,6^.

Studying invasion in the laboratory can be complicated due to the complexity of many microbial communities. For example, the human gut microbiome consists of hundreds of species, making it difficult to characterize specific microbe-microbe interactions^7,8^. *Drosophila melanogaster* is an attractive model for the study of microbial invasion for several reasons. Its intestinal tract is structurally and functionally similar to that of vertebrates, including humans, while having a more simplistic microbiome composition that can be controlled and manipulated in the laboratory^9–13^. Previous studies in *D. melanogaster* have examined microbial invasion through the lenses of microbial pathogenesis^14–18^, microbiome assembly^10,19^, and probiotic colonization^20,21^. In this study, we explore the effect of the gut microbiome on the invasion of three Gram-negative bacteria previously used in *D. melanogaster* studies: the lethal entomopathogen *Pseudomonas entomophila* (Pe), the non-lethal pathogen *Pectobacterium carotovora* (Ecc15), and the human probiotic organism *Escherichia coli* Nissle 1917 (EcN). Here, we report that the microbiome protects *D. melanogaster* during microbial invasion, leading to improved probability of survival and reduced microbial burden compared to axenic (microbiome-free) hosts. We report antimicrobial activity by the gut microbiome member *Lactiplantibacillus plantarum* (Lp) against invasive microbes, which is linked to its ability to acidify the surrounding environment.

Another microbiome member, *Acetobacter tropicalis* (At) reduces the inhibitory capacity of Lp by neutralizing acidity. Overall, this work expands our understanding of microbe-microbe interactions by characterizing acidity from the microbiome as a chemical barrier to microbial invasion, a phenomenon that can be observed in many animals, including humans^22–25^.

## Results

### The gut microbiome protects the host during invasion with a lethal pathogen

To determine how common members of the *D. melanogaster* microbiome impact host susceptibility to Pe infection, we generated axenic flies without microbiomes by hypochlorite dechorionation of embryos^26^. We then generated gnotobiotic animals by re-associating axenic flies with defined microbial communities: Lp, At, or LpAt (**Figure 1A**). During infection with Pe, each gnotobiotic treatment group experienced improved levels of survival compared to infected axenic flies (**Figure 1B**). Lp mono-colonization resulted in statistically better survival compared to mono-colonization with At. Co-colonization of the two gut microbes resulted in survival comparable to mono-colonization with Lp and At alone. We also assessed the microbial burden of Pe over the course of the infection and found that, while Pe load remained high over 7 days in axenic flies, the presence of Lp in the mono-colonized gnotobiotic condition resulted in lower levels of Pe compared to axenic flies at 72 hours and 7 days (**Figures 1C, S1)**. Pe levels in At mono-colonized flies remained high, similar to axenic conditions. When flies contained both Lp and At prior to infection, Pe load was reduced at 72 hours compared to axenic flies, but it increased again by 7 days. Together, these results demonstrate that the microbiome, particularly Lp, limits the colonization ability of Pe in the gut, which is consistent with the reduction in mortality observed in gnotobiotic flies.

**Figure 1.**
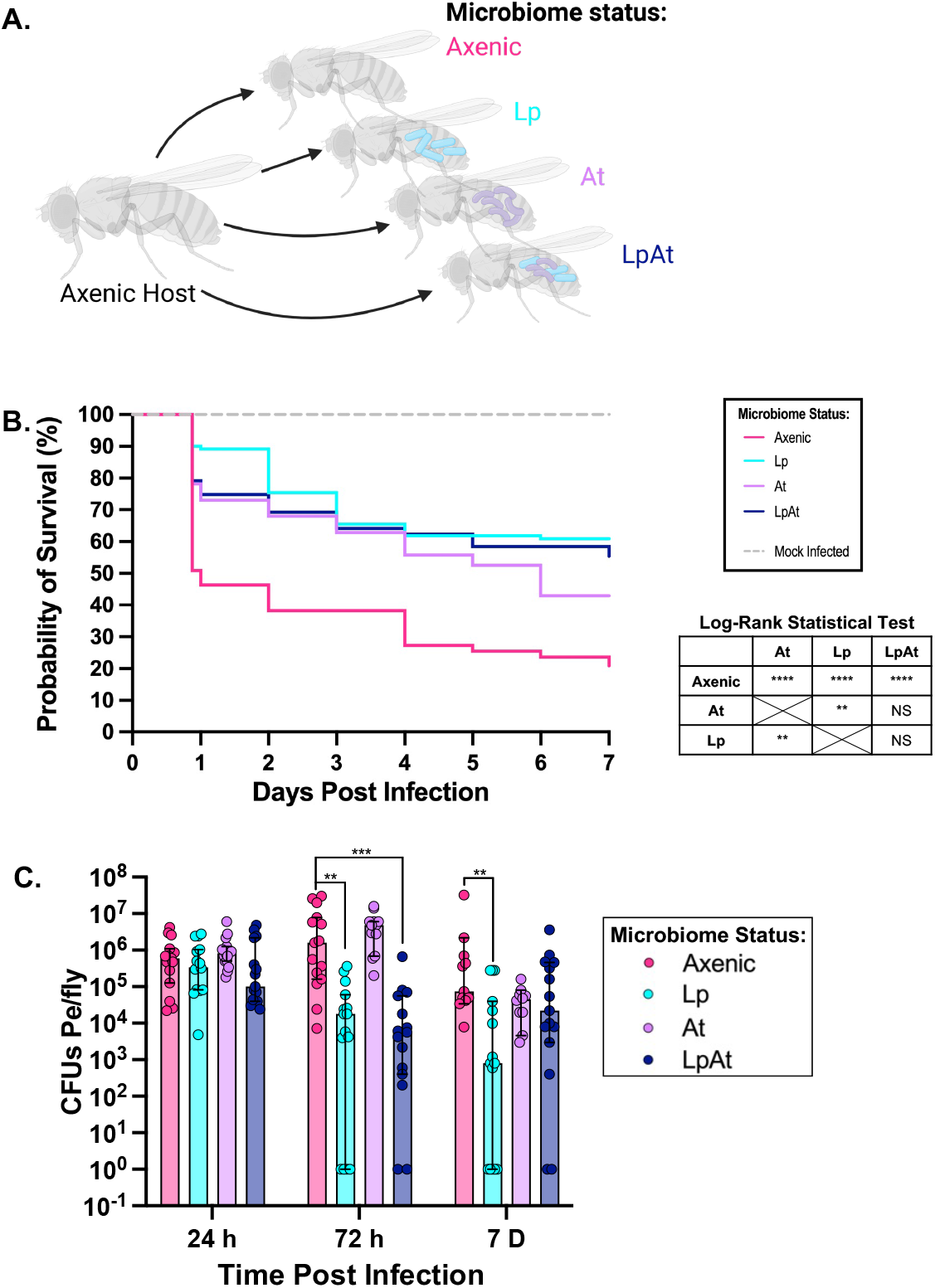
Presence of microbiome members reduces host susceptibility to *P. entomophila*. **A)** Scheme describing the generation of gnotobiotic flies. **B)** Kaplan-Meier survival analysis of female flies infected with *P. entomophila* (Pe) via feeding (feeding occurred from t=0 to t=1 day); fly microbiome treatments included no bacteria (Axenic), *L. plantarum* (Lp), *A. tropicalis* (At), or a combination of Lp and At (Lp/At). The mock infected group shows survival of axenic flies fed LB media; survival of all gnotobiotic conditions fed LB were also recorded but were not significantly different to axenic control and are not shown. Log-rank statistical analyses of each infection condition are compared to the other conditions. Significance is expressed as follows: NS, not significant; **, P ≤ 0.01; ****, P ≤ 0.0001. n=60 flies per control & 110-160 flies per Pe-infected treatment over 3 independent replicates. **C)** Number of colony forming units (CFUs) of bacteria per fly infected with Pe screened in axenic flies and gnotobiotic flies colonized with Lp, At, and Lp/At. Flies were sacrificed to determine Pe load at 24 hours, 72 hours, and 7 days post feeding infection. Each point represents bacterial load from an individual fly; bars and error bars represent the median and 95% confidence intervals; limit of detection is 2×10^2^ CFUs/Fly. n=9 flies per control & 10-15 flies (depending on availability of living flies) per infected treatment per time point over 3 biological replicates. Statistical significance was determined between flies containing different microbiomes by the Kruskal-Wallis method with Dunn’s multiple comparisons analysis. P-values are represented as follows: *, P≤0.05; **, P≤0.01; ***, P≤ 0.001; ****, P≤ 0.0001.

### Microbial invasion by Gram-negative microbes is reduced in flies associated with *L. plantarum*

Previous studies in *D. melanogaster* have demonstrated that the microbiome protects the host during challenge with lethal pathogens^27^. However, the role of the microbiome during the introduction of non-lethal invasive microbes has largely gone unaddressed. We selected two such Gram-negative bacteria: *Pectobacterium carotovora* (Ecc15), a plant pathogen that has adapted to cause gut disease in flies^15^, and *Escherichia coli* Nissle 1917 (EcN), which we previously identified as a colonization proficient strain of *E. coli* that does not cause obvious disease in flies^20^. We administered both organisms to axenic and gnotobiotic flies and assessed microbial burden over the following 48 hour period to probe the impacts of the microbiome on the outcome of infection. We found that the Ecc15 microbial load was reduced in all three microbiome treatments compared to axenic flies at 24 hours post-infection, with Lp-associated flies exhibiting the greatest diminution (**Figure 2A**). Surprisingly, EcN bacterial load was not greatly affected by the presence of Lp, but slight differences in bacterial load were observed at 24 hours between axenic and At-gnotobiotes and at 48 hours between axenic and gnotobiotes harboring At alone or the combination of Lp and At **(Figure 2B)**. Taken together with the Pe infection data, these results strongly suggest that the presence of Lp protected the host during invasion of pathogenic microbes, and we sought to uncover the mechanism of this protection.

**Figure 2.**
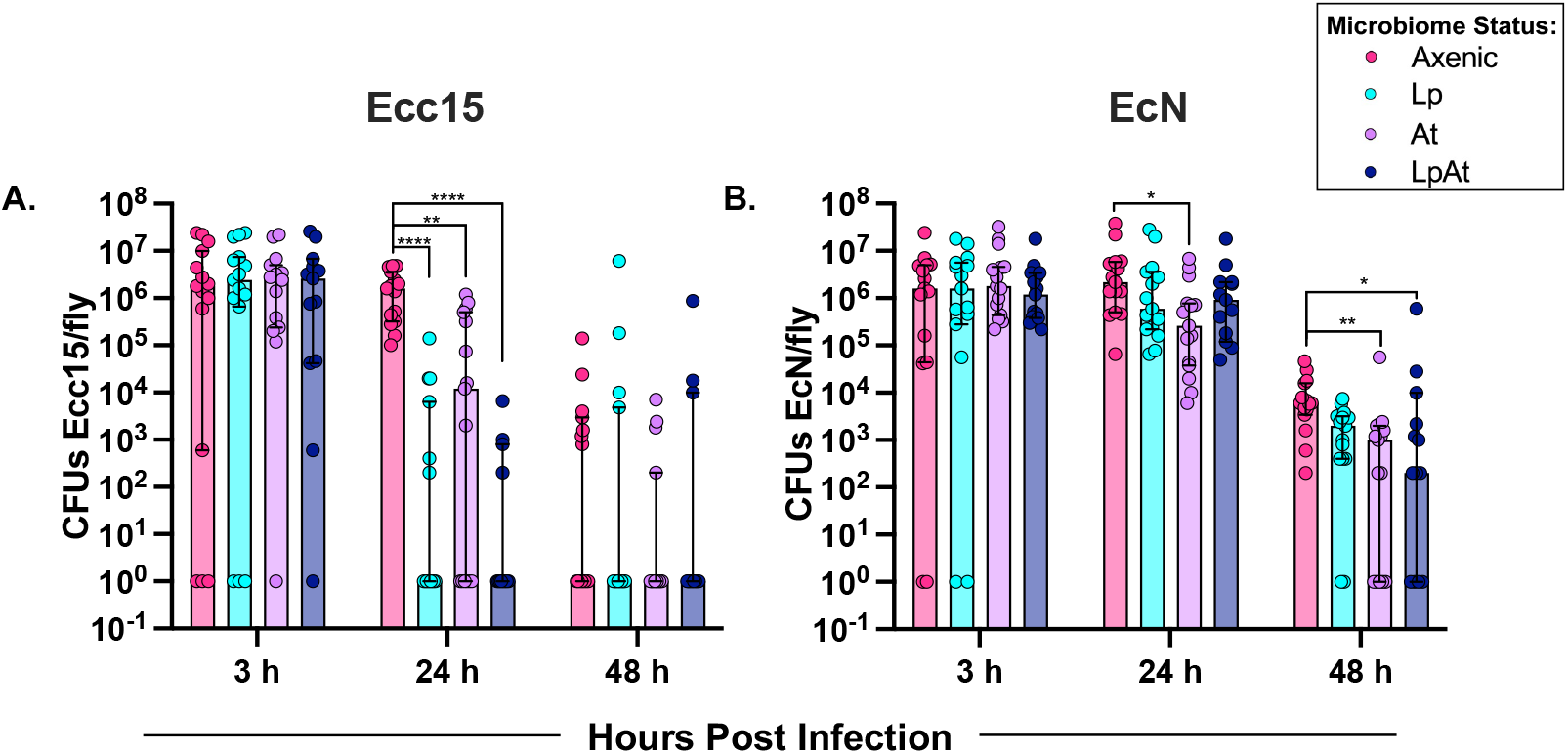
Microbiome composition alters the microbial load of invasive organisms during infection. Number of colony forming units (CFUs) of bacteria per fly infected with Ecc15 (**A**), and EcN (**B**). Each invasive organism was screened in axenic flies and gnotobiotic flies colonized with Lp, At, and Lp/At. Bacterial load was determined at 3 hours, 24 hours, and 48 hours post feeding infection. Each point represents bacterial load from an individual fly; bars and error bars represent the median and 95% confidence intervals; limit of detection is 2×10^2^ CFUs/Fly. n=15 flies per treatment per time point over 3 biological replicates. Statistical significance was determined between flies containing different microbiomes by the Kruskal-Wallis method with Dunn’s multiple comparisons analysis. P-values are represented as follows: *, P≤0.05; **, P≤0.01; ***, P≤ 0.001; ****, P≤ 0.0001.

### *L. plantarum* inhibits invasive organisms *in vitro*

Traditionally, the “cross-streak” assay has been used as an *in vitro* culture-based method to study binary microbe-microbe interactions. To expand this method to examine the combined effects of two microbiome organisms on the growth of invasive microbes, we developed the multi-organism interactions assay **(Figure 3A)**. In this assay, intersecting lines of Lp and At were spread at a 90° angle in the middle of a Petri dish and allowed to grow for three days, giving enough time for the microbes to secrete metabolites into the environment (**Figure S2**). The invasive organism (Pe, Ecc15, or EcN) was then streaked into the adjacent quadrants and grown for an additional 1-2 days. These assays revealed that Lp, but not At, inhibited the growth of Pe, Ecc15, and EcN, as evidenced by a zone of inhibition near the Lp streak and growth close to the At streak (**Figure 3B,** see dashed regions). Interestingly, our assay also captured that At modulated the inhibitory effects of Lp on all three organisms, as the zones of inhibition diminish away from the distal end of the Lp streak, practically disappearing at the intersection point of Lp and At (**Figure 3B**). We observed the same patterns of inhibition when Pe, Ecc15, and EcN were co-cultured with fly microbiome isolates in liquid media. Co-culturing with At did not impact growth of Pe, Ecc15, or EcN, co-culturing with Lp alone inhibited their growth, and the addition of At reduced this inhibition by Lp (**Figure 3C**). Interestingly, EcN was more resilient during co-culture with Lp than Pe and Ecc15, which were completely inhibited, but populations were still ~1000x less abundant than when grown on its own. Altogether, these results demonstrate that Lp has an antimicrobial effect on the growth of the three invasive organisms, which At can ameliorate by creating an environment more favorable for their growth.

**Figure 3.**
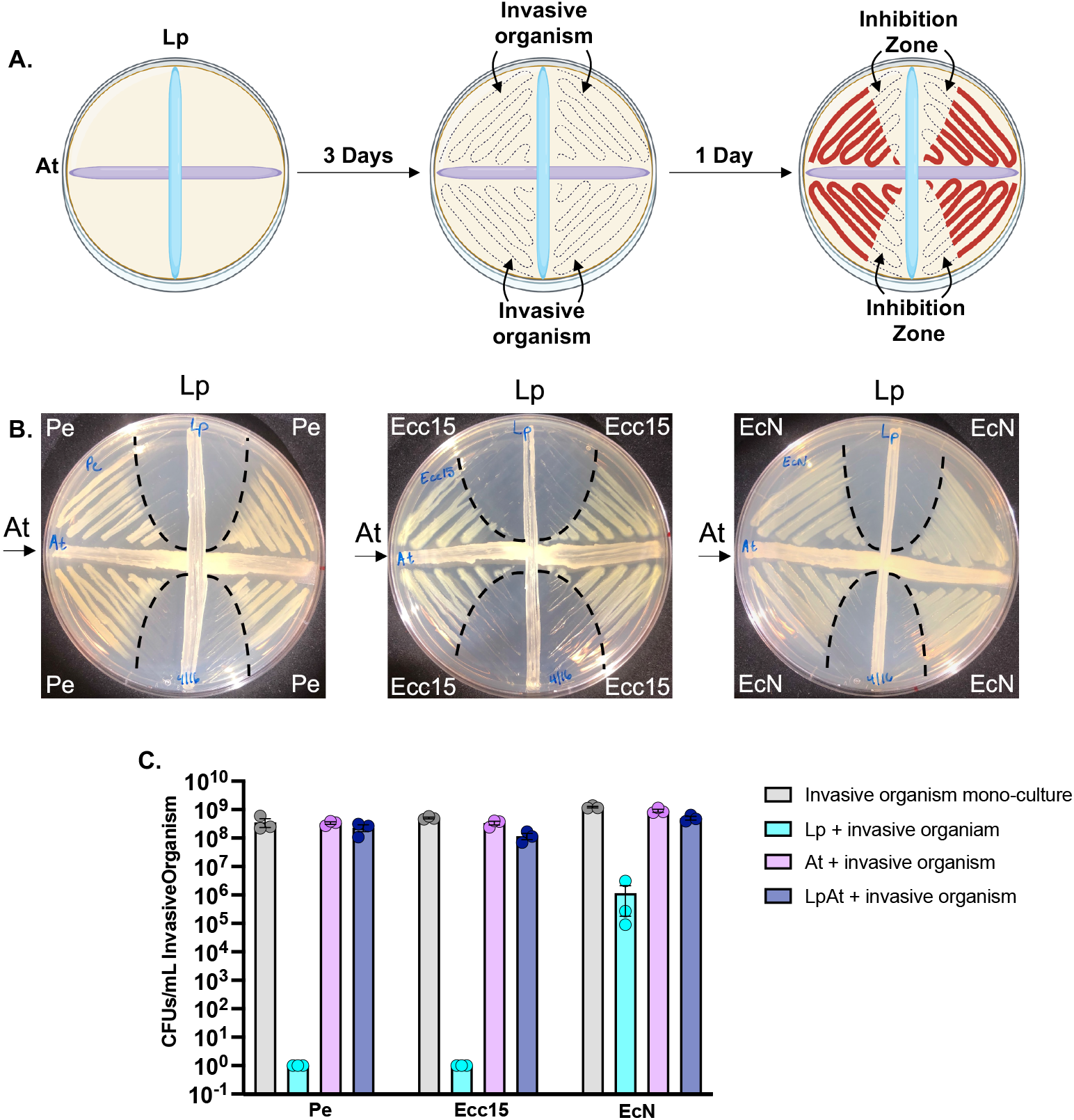
*In vitro* growth assays reveal inhibition of Gram-negative invasive organisms by Lp. **A)** Scheme describing the multi-organism interaction assay procedure. **B)** Multi-organism interaction assays display growth effects of gut microbes Lp and At on invasive organisms Pe, Ecc15, and EcN. Microbiome members were grown in perpendicular streaks for 3 days, and invasive organisms were added to the adjacent quadrants and allowed to grow for an additional day. Zones of inhibition are indicated by dashed lines. **C)** Co-culture analysis of invasive organisms with microbiome members. Pe, Ecc15, and EcN were grown in mono-culture or in media previously inoculated with Lp, At, or a mixture of both (LpAt). Bars and error bars represent the mean concentration of invasive organisms in CFUs/mL ±SEM. N=3 biological replicates per condition.

### *L. plantarum-mediated* inhibition is driven by environmental acidification

Lactic acid (and its conjugate base, lactate), a major Lp metabolite, has recently been implicated as an important factor in microbe-microbe interactions and microbe-host interactions in *D. melanogaster*^28–30^. To determine if the secretion of lactic acid is a potential mechanism by which Lp inhibits the growth of the selected invasive organisms, we measured the pH of the media surrounding Lp in various growth conditions. We first observed the pH of media in which Lp and At were grown at the same time (streaked perpendicular to one another as in **Figure 3A**) using media plates containing the pH indicator bromophenol blue, and saw distinct acidic regions (as indicated by a yellow color) surrounding Lp, with larger acid penetrance further away from the interaction point with At (**Figure 4A**). Introduction of the invasive organisms yielded zones of inhibition overlapping with the zones of acidity (**Figures 4A, S3**). Interestingly, while Pe was inhibited as in **Figure 3B**, the acidic zone surrounding Lp returned to a blue color, indicating that the pH had risen back above 4.6 (**Figure S3**). This experiment suggested that the ability of Lp to acidify the surrounding media is linked to its inhibition of invasive microbes and that At reduces its inhibitory capacity by neutralizing acidity. To mimic the growth conditions of the microbiome members *in vitro,* we determined the pH of overnight cultures of Lp and At mono-cultures and Lp/At co-cultures (**Figure 4B**). The Lp culture was found to be pH 4.0, consistent with its known capacity to secrete lactic acid. *A. tropicalis* had a pH of 7.0, and the Lp/At co-culture had a pH of 6.0. To test the effect of low pH on the growth of the invasive microbes, culture media was adjusted to pH 4.0 with either lactic acid or hydrochloric acid. Invasive microbes were grown alone in standard media and acidified media and in co-culture with At in acidified media. The lactic acid media completely prevented the growth of the Pe, Ecc15, and EcN (**Figure 4C**). Media acidified with HCl reduced (but did not eliminate) the culture densities of Pe and Ecc15, while having little effect on the growth of EcN (**Figure S4)**. Co-culturing with At in both acidified media treatments restored the growth of each organism to normal levels, while also raising the pH of each culture to >5.0 (data not shown). To complement this analysis, we co-cultured each invasive organism with Lp in standard media (starting pH 6.5) and media buffered to pH 6.0 with 60 mM phosphate buffer (**Figure 4D**). As in **Figure 3C**, Pe and Ecc15 were completely inhibited by Lp, and EcN was partially inhibited in standard media. However, all three organisms grew to a normal level when co-cultured with Lp in buffered media, suggesting that Lp-mediated inhibition was linked to its ability to shift the pH of the surrounding environment. To test this hypothesis *in vivo,* we performed bacterial load analysis on axenic and gnotobiotic flies infected with Ecc15 on the standard fly diet and on a fly diet buffered to pH 6.0 with 2-(*M*-morpholino)ethanesulfonic acid (MES) (**Figures 4E, S5**). Interestingly, axenic and Lp-colonized flies feeding on the buffered diet had significantly higher levels of Ecc15 in their intestines 48 hours-post infection. This demonstrates that acidification of the gut is crucial to the protective effect of Lp during infection. To assess gut pH, we fed axenic and gnotobiotic flies bromophenol blue for 24 hours, dissected their guts, and imaged them to assess intestinal pH (**Figure 4F**). We found that flies colonized with either Lp or At had distinct acidic copper cell regions (indicated by a green-yellow color) while the guts of axenic flies or flies colonized with Lp and At lacked such distinct regions, suggesting that microbiome composition alters the pH of the intestine.

**Figure 4.**
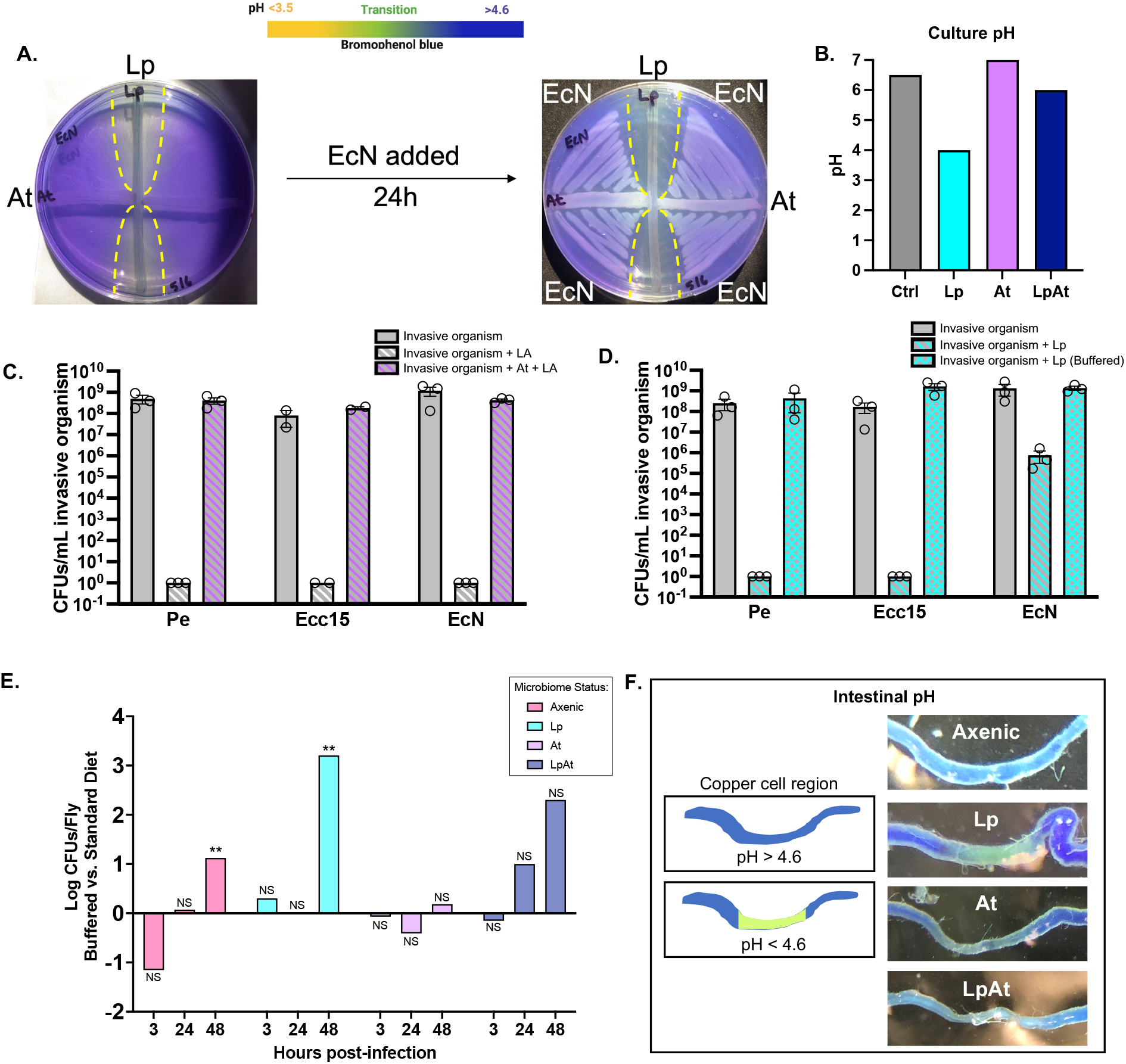
Microbiome-derived shifts in pH contribute to inhibition by Lp. **A)** Multi-organism interaction assay showing the effects of Lp and At on EcN growth on media containing the pH indicator bromophenol blue. After three days of growth, a yellow acidic zone appears around Lp, but it tapers off near the At interaction point. When EcN is added, the zone of inhibition overlaps with the acidic region. **B)** pH measurement of microbiome mono-cultures and co-cultures reveals a sharp acidification of culture media by Lp. **C)** Density of invasive organisms in media adjusted to pH 4.0 with lactic acid (LA) with or without 24 hours of prior growth with At. Each bar represents mean CFUs/mL ±SEM of 3 biological replicates. **D)** Microbial concentration of invasive organisms in media buffered to pH 6.0 with phosphate buffer with or without 24 hours of prior growth with Lp. Each bar represents mean CFUs/mL ±SEM for 3 biological replicates. **E)** Ratio of Ecc15 microbial load in flies infected on buffered vs. standard fly diets, n=15 flies per treatment per time point over three biological replicates. Statistical significance was determined for flies of each microbiome status between standard and buffered diets using the Kruskal-Wallis method with Dunn’s multiple comparisons analysis. P-values are represented as follows: NS, P>0.05; **, P ≤ 0.01. **F)** The pH of the copper cell region of the intestine was approximated by feeding axenic and gnotobiotic flies food soaked with 2% bromophenol blue. Guts were dissected and imaged immediately. A yellow/green color in the copper cell region indicates an acidified environment (pH < 4.6).

### Microbiome composition alters the chemical environment of fly food

The observation that Lp rapidly acidifies bacterial culture media raised a question about the effect of the microbiome members on the chemistry of fly food. Fly food is a rich source of microbes as flies feed upon and defecate microbes continuously, and as such, it has been described as a “reservoir” for the microbiome^27^. We tested the effects of gut microbes on the pH of fly food in two different ways: by applying bacterial cultures directly to fly food (**Figure 5A**) and by introducing axenic and gnotobiotic flies to vials of fly food (**Figure 5B**). In both experiments, the control fly food was ~pH 6.0. Fly food colonized by Lp was reduced to ~pH 4.0. *A. tropicalis* had more minimal impacts on food pH, with slightly higher pH (6.5) when culture-inoculated and slightly lower pH (5.5) when fly-inoculated. Interestingly, the pH of fly food co-inoculated with Lp and At was ~pH 5.0 in both experiments, indicating a level of acidity intermediate between Lp and At alone. These data demonstrate that microbiome composition has a critical impact on the pH of fly food, as fly food containing Lp alone was found to be ~100x more acidic than uninoculated controls, while At had a less dramatic effect on pH. Given the different levels of acidity observed in the presence of different gut microbes, we asked whether the invasive organisms experience similar interactions with gut microbes on fly food. To test this, we added gut microbes to the surface of fly food, allowed them to grow for three days, and applied Pe, Ecc15, or EcN to the same vial and cultured them from the fly food 24 hours later to determine the levels of invasive microbes and microbiome members (**Figure 5C, Figure S6)**. As observed in previous analyses, At had little effect on the growth of the invasive microbes on fly food. Lp completely prevented the growth of Pe and Ecc15, while greatly reducing the level of EcN growth compared to the control. Levels of invasive microbes were partially rescued when Lp and At were grown together on fly food, suggesting that At alters the inhibitory capacity of Lp on fly food. These experiments support the idea that fly food acts as a reservoir for not only the microbiome community members, but also the metabolic products they release into the environment. These include acidic products such as lactic acid, which have the potential to play an important role in modulating the environment and affecting both host and microbial processes **(Figure 5D)**.

**Figure 5.**
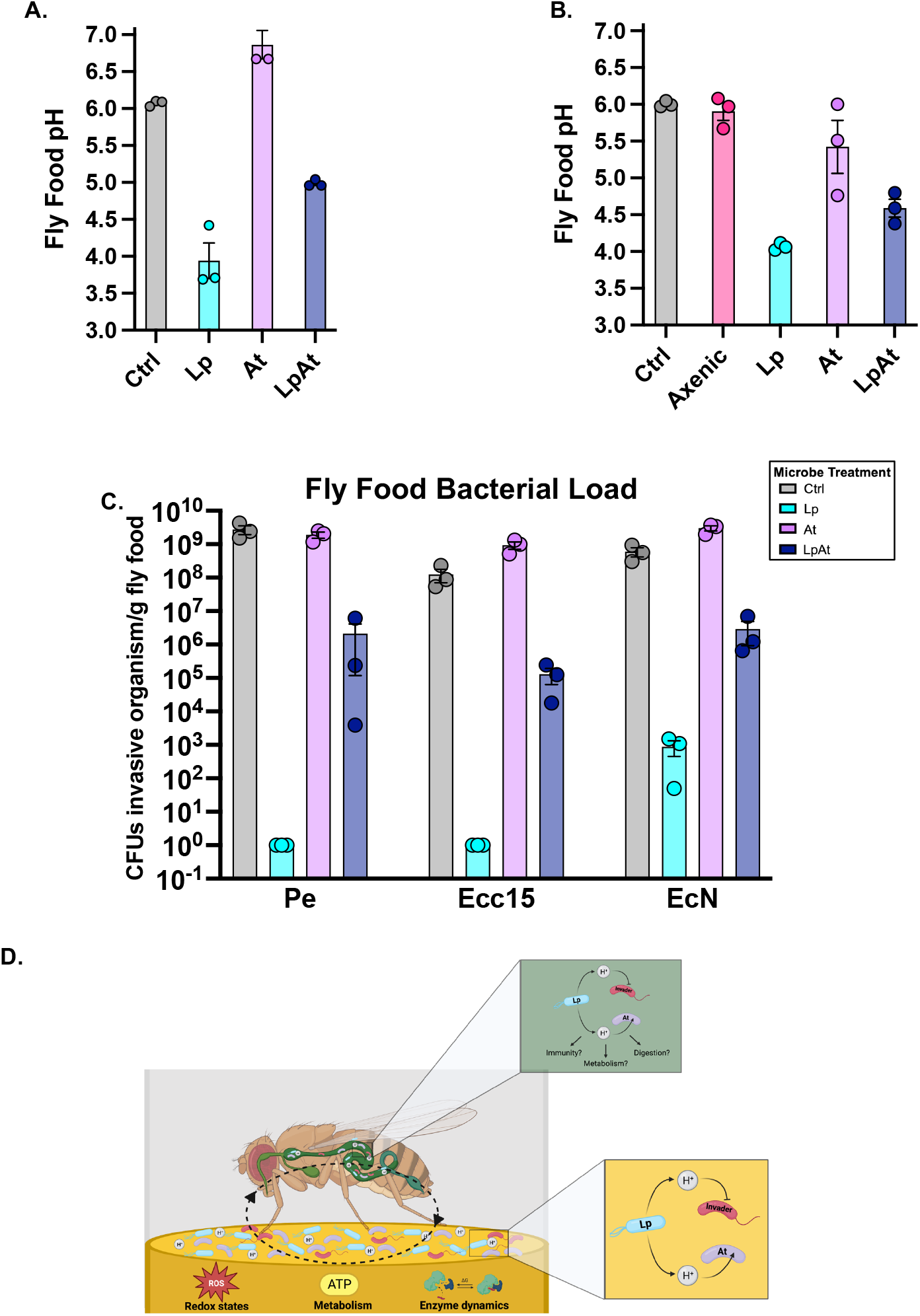
Microbiome composition alters the chemical environment of fly food. **A)** pH analysis of fly food after three days of growth of Lp, At, or Lp/At, with culture media added as a control. Bars represent the mean pH of three biological replicates ±SEM. **B)** pH analysis of fly food three days after the addition of 40 male flies of different microbiome statuses (Axenic, Lp, At, Lp/At). Bars represent the mean pH ±SEM of three biological replicates. **C)** Bacterial load of Pe, Ecc15, and EcN present on fly food initially inoculated with microbiome members Lp, At, or Lp/At, with culture media as a control. Each bar represents the mean CFUs/g fly food ±SEM for three biological replicates. **D)** Pictorial representation of microbe-microbe interactions between microbiome members and Gram-negative invasive bacteria on fly food and during consumption by the host.

## Discussion

### The microbiome protects the host during infection with invasive organisms

We examined microbe-microbe interactions in a *D. melanogaster* model, which has the advantage of possessing a simple microbial community that can be removed and manipulated to provide a high level of control over the system. Previous work showed that axenic flies had reduced survival during infection with *Pseudomonas aeruginosa* and *Serratia marcescens* compared to flies raised with their microbiomes intact^27^. Our study expanded upon this finding by assessing survival and microbial burden in gnotobiotic flies with defined microbial communities. We found that flies harboring gut microbes *Lactiplantibacillus plantarum* (Lp), *Acetobacter tropicalis* (At), or a consortium of the two had an improved probability of survival compared to axenic flies when infected with the gut pathogen *Pseudomonas entomophila* (Pe) and that this was accompanied by a reduction in microbial burden of Pe, most starkly in the Lp gnotobiotes (**Figure 1**). We questioned whether the apparent protective effect of the microbiome was specific to lethal pathogens or whether a more generalized mechanism of protection was at play. To test this, we screened two other microbiome-invasive but non-lethal microbes *Pectobacterium carotovora* (Ecc15) and *Escherichia coli Nissle 1917* (EcN) in our model and observed a similar reduction in microbial burden of Ecc15 in Lp gnotobiotes, but no such reduction of EcN, suggesting that EcN is a better invader into the microbiome (**Figure 2**).

### Weak organic acids act as gatekeepers for invasion of microbial communities

Microbial invasion has been studied extensively and has implications on a variety of biological processes from pathogenesis to agriculture. It involves the introduction and establishment of a microorganism, initially foreign, into a new environment^1^. Multiple biotic and abiotic barriers must be overcome before a microbe can successfully invade an existing microbial community^1^. In this study, we characterized microbiome-derived acidity as a chemical barrier to microbial invasion. *In vitro* analyses of microbe-microbe interactions revealed that Lp inhibited the three invasive organisms we tested, and the observation that At lessened this inhibition when grown in close proximity to Lp was a key finding as it suggested that At modulates the environment to make it more favorable for invasive organisms (**Figure 3**). We found a link between the microbial inhibition phenotype and the pH of the culture media, as evidenced by the acidification of the surrounding environment by Lp and neutralization of acidity by At (**Figure 4**). These findings suggest that the composition and invasion resiliency of the microbiome is tightly linked to the biochemical environment created by the resident microbes. We hypothesize that acidity from the microbiome acts as a chemical barrier and gatekeeper to the establishment of an invader. While this harsh environment eliminates many potential invaders, microbes that overcome this hurdle can become established and eventually displace the resident community. Another interesting finding from our study was that not all acids have the same inhibitory capacities, even when set at an identical pH. For example, when we adjusted culture media to pH 4.0, lactic acid completely inhibited EcN, while hydrochloric acid had little effect on its growth (**Figures 4C, S4**). A similar finding was made with a human isolate of *E. coli* O157:H7^31^. This is likely explained by the ability of the acid to enter the microbial cytoplasm. Strong acids such as HCl (pK_a_ −5.1) fully dissociate in aqueous environments, and the positively charged H^+^ ions cannot readily permeate the plasma membrane^32,33^. This contrasts with weak acids such as lactic acid (pK_a_ 3.86), which do not fully dissociate and can diffuse more easily into the microbial cell where they can dump their protons and acidify the cytoplasm. This causes issues with protein stability, redox balance, and the proton motive force^32^ (**Figure S4**). Considering the types of acids present and their abundance, and how different microbiome compositions shape these attributes will be of interest for future study. For example, in our system the amount of lactic acid needed to match the Lp culture pH (~4.0) was 3.5 times higher than the amount of lactate we measured in these cultures, suggesting additional acids are contributing to the final pH and inhibition by Lp (**Figure S4**).

### Establishing common rules for microbial invasion across hosts

Taken together, our results add to the growing body of evidence describing the rules for the control of microbial invasion. The deployment of acids acts as an initial sieve to block the establishment of foreign invaders, with later, more specific interventions by the host immune response and direct competition by the microbiome also acting as barriers^1,6,13,16,34,35^. The finding that Lp protects the host by acidifying the environment parallels interactions observed in the human vaginal microbiome. In particular, *Lactobacillus crispatus* plays an important role in maintaining homeostatic conditions by producing weak organic acids that maintain the pH of the lower reproductive tracts of reproductive-aged women at ~ 4.5^22,36^. If an invasive organism perturbs the vaginal microbial community and neutralizes the acidity (similarly to At in our model), the host becomes more susceptible to opportunistic pathogens such as *Candida albicans*^25,37^. This is consistent with our findings and bolsters the utility of the *D. melanogaster* gut as a model, as it serves as a vacuum to study inter-species microbial interactions on epithelial surfaces in granular detail.

### Fly food as a reservoir for the microbiome and its metabolites

When considering how the *D. melanogaster* microbiome interacts with external microbes encountered by the host, the microenvironment associated with fly food is an important consideration. Fly food has previously been suggested to behave as a “reservoir” for the microbiome, as frequent transfer of flies to sterile fly food diminishes the microbiome load observed^27^. We previously established that fly microbiome community members alter the nutritional content of fly food, leading to an increased protein-to-carbohydrate ratio in the diet^38,39^. In the present study, we have expanded our understanding of how the microbiome affects the chemical makeup of fly food by demonstrating strong acidification of the diet by Lp, which is absent when At alone is added to the food. Interestingly, when Lp and At are both added to fly food, the level of acidity is intermediate, suggesting that At quenches some of the acidic compounds produced by Lp, which is unsurprising considering that many *Acetobacter* species utilize lactate as a carbon source^40,41^. This is an important observation because quite often, the fly food environment is overlooked as a contributor to host physiology. We argue that under laboratory conditions, as in wild *D. melanogaster* populations, the food substrate is a critical component of the host-microbe relationship in that it acts as a source and a sink for the microbiome, as well as its metabolites. Moreover, these metabolites and biochemical impacts likely alter food even prior to fly ingestion (**Figure 5D**). Acidic fly diets were previously shown to increase fly gustatory responses and food intake while also extending lifespan^42^. Another study found that interactions among microbiome members leading to the production of acetic acid protected developing larvae against pathogenic fungi and influenced host behavior^43^. These interactions and their effects on the chemical environment could potentially impact invasive microbes by altering redox states, metabolites, and enzyme kinetics. Inside the host gut environment, the microbiome and its acidic products may play direct roles in modulating immunity, metabolism, and digestion^34,44,45^. Going forward, this will be an important consideration when designing host-microbe studies in *D. melanogaster* as the microbiome’s relationship with the fly diet comes further into focus.

## Materials and Methods

### Fly stocks and rearing

Oregon-R flies obtained from the Bloomington *Drosophila* Stock Center (BDSC #5) were initially treated with a final concentration of 0.05% tetracycline in food to eliminate *Wolbachia* and were maintained for at least an additional three generations prior to starting experiments. Stocks were maintained at 25°C with 12 hour light:dark cycling on the Broderick Standard diet^46^ containing, per liter: 50 grams inactive dry yeast, 70 grams yellow cornmeal, 40 grams sucrose, 6 grams *Drosophila* agar, and 1.25 grams methyl paraben (dissolved first in 5 mL of 100% ethanol). Fly food was autoclaved before use and newly emerged adults were passaged into new tubes every 3-4 days.

### Bacterial culturing methods

Unless otherwise specified, bacterial cultures were prepared as follows:

| Organism | Media | T°C | Reference |
| --- | --- | --- | --- |
| *Lactiplantibacillus plantarum* (Lp) | MRS | 29 | Broderick et al., 2014 <sup>26</sup> |
| *Acetobacter tropicalis* (At) | MRS | 29 | Judd et al., 2018 <sup>47</sup> |
| *Pseudomonas entomophila* (Pe) | LB | 29 | Vodovar et al., 2005 <sup>14</sup> |
| *Pectobacterium carotovora* (Ecc15) | LB | 29 | Basset et al., 2003 <sup>15</sup> |
| *Escherichia coli* Nissle 1917 (EcN) | LB | 37 | Mutaflor ® |

*Pseudomonas entomophila* (Pe). GFP-expressing *Pseudomonas entomophila* was grown from −80°C on Lysogeny broth (LB-10g tryptone, 5g yeast extract, 5g NaCl) agar containing 1% milk and incubated at 29°C for two days. Colonies expressing GFP and protease activity (visualized as clearing in milk plates) were grown shaking in liquid LB containing 3% NaCl for 20 hours at 29°C.

### Generation of axenic and gnotobiotic flies

Adult flies were kept in an embryo collection chamber with grape juice agar smeared with yeast paste overnight. Embryos were rinsed with ethanol, dissociated from the grape juice agar using PBS and sterile swabs, and collected in a cell strainer. The chorion was removed using 5% bleach and dechorionated embryos were rinsed with sterile water and ethanol before being transferred via pipette to sterile medium. Vials of food were exposed to ultra-violet light for 15 minutes before use for axenic flies, and all flipping of axenic stocks was done in a biosafety cabinet or near a flame.

To generate gnotobiotic flies, *L. plantarum* and *A. tropicalis* were grown, overnight cultures were set to OD_600nm_ = 0.5, and 150 μL of the diluted cultures (or a 1:1 mix of both) were fed to axenic flies. Stocks of gnotobiotes were maintained for at least six generations prior to being used in infection experiments.

### Preparing invasive organisms for oral infection

Liquid cultures of Pe, Ecc15, and EcN were centrifuged for 20 minutes at 3700 rpm at 4°C, the supernatant was removed, and the pellets were resuspended by light vortexing. Cultures were adjusted using media to OD_600nm_=200 (OD200) for infections. Bacterial pellets (or media for controls) were mixed in a 1:1 ratio with 2.5% sucrose, and 140 μL of this infection medium was used to saturate paper filters on top of fly food.

Adult female flies (4-7 days old) were starved for 2 hours at 29°C before being transferred to infection tubes and kept at 29°C for the duration of the experiment. Pathogen-independent deaths were recorded at t=2 hours. Experimental vials were passaged at 24 hours, then every two days (for Pe).

### Bacterial load analysis

For *P. entomophila* infection analysis, two sets of flies were infected as described above per replicate (N=3 replicate infections); one set per replicate infection was monitored for survival while individuals were taken from the other set for assessment of bacterial load of Pe, Lp, and At at 24 hours, 72 hours, and 7 days post-infection start. Flies infected with Ecc15 and EcN were sacrificed at 3 hours, 24 hours, and 48 hours post-infection start and assessed for bacterial load. Each infection vial was subjected to CO2 anesthetization, and a sub-sample of individuals was removed, surface sterilized in 70% ethanol, rinsed in sterile PBS, and homogenized in sterile screw-top vials containing glass beads and 0.6 mL of PBS. 3-5 flies were taken at each time point from LB control infection tubes, and five flies were taken from invasive organism-infected tubes (or as many living flies were left, up to five). Following homogenization, samples were diluted to 10^-5^ and spot-plated as 3 μL spots. Pe-containing samples were plated on LB+milk agar. Ecc15- and EcN-containing samples were plated on LB agar. All samples were also plated on MRS agar to capture Lp and At. Samples containing EcN were also plated on MRS agar supplemented with 50 μg/ml ampicillin. All plates were incubated at 29°C. Counts were recorded 1-2 days later.

### Visualization of *in vitro* microbe-microbe interactions

#### Multi-organism interaction assays

To visualize the combined effects of Lp-At interactions on the growth of invasive organisms, Lp and At were grown in perpendicular lines on a Petri dish containing mannitol agar for two days. Each invasive organism was streaked in the adjacent quadrants and grown for an additional day. Plates were photographed, and inhibition of the test organism was measured in millimeters proximal and distal from the point of interaction between Lp and At.

#### In vitro pH approximation

pH of interactions on plates was approximated by supplementing mannitol agar with 0.02% bromophenol blue, which turns yellow at ~pH 3.5. The pH of bacterial cultures was determined using universal pH indicator solution (Sigma).

#### Co-culture assays

Lp and At were grown in MRS broth for 18 hours, shaking at 200 rpm at 29°C. Both cultures were adjusted to OD_600_=0.2 (OD0.2). These starter cultures were used to inoculate experimental co-cultures by spiking 100 μl of Lp, At, or a 1:1 mix into test tubes containing 5 mL of mannitol broth. Mono-cultures of Pe, Ecc15, and EcN were prepared in 5 mL of LB. Mono-cultures and co-cultures were grown for 18 hours, shaking at 200 rpm at 29°C. Mono-cultures of invasive organisms were adjusted to OD0.2, and 100 μl of the appropriate organism was spiked into each co-culture. Mono-cultures of the invasive organisms were also prepared in 5 mL of mannitol broth from the OD0.2 starter cultures for comparison. All cultures were grown for an additional 18 hours, shaking at 200 rpm at 29°C. Cultures were serially diluted and plated on both LB agar and MRS agar and grown for 1-2 days at 29°C.

#### Lactic acid determination

Lactic acid concentrations of overnight cultures of Lp, At, and LpAt co-cultures were determined using the Lactate Assay Kit II (Sigma) according to the manufacturers’ instructions.

#### Acid co-cultures

To mimic the lactic acid content of an Lp culture, mannitol media was supplemented with 4 mM lactic acid. To mimic the pH of an Lp culture, mannitol media was titrated with either lactic acid or hydrochloric acid to pH 4.0. *A. tropicalis* was grown for 18 hours in MRS broth, shaking at 200 rpm at 29°C and adjusted to OD_600_=0.2. Two sets of cultures were set up and grown overnight in test tubes containing 5 mL of acid-supplemented media: one set spiked with 100μl of At and one set lacking At. Pe, Ecc15, and EcN were grown overnight in LB, adjusted to OD_600_=0.2, and 100 μl was spiked into the appropriate acid-supplemented media tube. The co-cultures were grown for an additional 18 hours. Cultures were serially diluted and plated on both LB agar and MRS agar and grown for 1-2 days at 29°C.

#### Buffered co-cultures

Mannitol media was supplemented with 60 mM phosphate buffer (Na(HPO_4_)_2_ + Na_2_HPO_4_). The media was titrated with HCl to pH 6.0. Lp was grown overnight and adjusted to OD_600_=0.2. Two sets of Lp cultures were set up and grown overnight in test tubes containing 5 mL of either buffered or non-buffered mannitol media. Pe, Ecc15, and EcN were grown overnight in LB, adjusted to OD_600_=0.2, and 100 μl was spiked into the appropriate media tube. The co-cultures were grown for an additional 18 hours. Cultures were then serially diluted and plated to determine bacterial density.

#### Gut pH measurement

To approximate pH regionality of the gut, axenic and gnotobiotic flies were placed in fly food vials supplemented with 2% bromophenol blue indicator (Sigma). Flies were allowed to feed for 24 hours before being anesthetized and dissected. Intestines were rapidly photographed as gut contents quickly escape following dissection.

#### Buffered fly food infections

Fly food was prepared using the standard Broderick Lab recipe (see above) with the liquid portion supplemented with 60 mM 2-(*N*-morpholino)ethanesulfonic acid (MES). The resulting slurry was adjusted to pH 6.0 with NaOH, distributed into vials, and autoclaved. Ecc15 infections were then carried out as described above.

#### Fly food pH measurement

For culture-inoculated fly food analysis, Lp and At were grown overnight in MRS media and adjusted to OD_600_=0.5. 150 μl of Lp, At, or a 1:1 mixture of Lp and At was applied to the surface of fly food and allowed to soak in for 3 days at 25°C. For gnotobiotic fly-inoculated fly food analysis, 40 axenic/gnotobiotic male flies were sorted aseptically, transferred to sterile fly food, and incubated for 3 days at 25°C. Fly food was removed from the vial and weighed, vortexed vigorously with deionized water, and the pH was measured using an Orion Star pH meter (ThermoFisher).

#### Fly food bacterial load determination

Lp and At were grown overnight and adjusted to OD_600_=0.5 with MRS. 150 μl of Lp, At, or an LpAt mix (~10^7^ cells total) were deposited on the surface of fly food vials, with MRS added as a control. After three days at 25°C, 150 μl of OD0.5 Pe, Ecc15, or EcN was added to the appropriate tubes, with LB as a control. 24 hours later, 1 g of fly food was aseptically collected, serially diluted, and plated on LB and MRS agar to determine the density of gut microbes and invasive organisms.

## Acknowledgements

The authors thank Dr. Arne Schon (Johns Hopkins Department of Biology) for his helpful discussions about the project. Figures were created with GraphPad Prism and BioRender.

This project was supported by NIH grant R35GM128871.

The authors have no conflicts of interest to declare.

A.J.B, D.N.A.L., J.D., and S.C. performed research, and A.J.B., D.N.A.L., and N.A.B. contributed to designing research, analyzing data, and writing the paper.

## References

1. Mallon, C. A., Dirk Van Elsas, J. & Salles, J. F. Microbial Invasions: The Process, Patterns, and Mechanisms. Trends Microbiol 23, 719–729 (2015).

2. Shade, A. et al. Fundamentals of microbial community resistance and resilience. Front Microbiol 3, 417 (2012).

3. Casadevall, A. & Pirofski, L. A. The damage-response framework of microbial pathogenesis. Nature Reviews Microbiology 2003 1:1 1, 17–24 (2003).

4. Gupta, S., Allen-Vercoe, E. & Petrof, E. O. Fecal microbiota transplantation: in perspective. Therap Adv Gastroenterol 9, 229 (2016).

5. Kaper, J. B., Nataro, J. P. & Mobley, H. L. T. Pathogenic Escherichia coli. Nature Reviews Microbiology 2004 2:2 2, 123–140 (2004).

6. Vila, J. C. C., Jones, M. L., Patel, M., Bell, T. & Rosindell, J. Uncovering the rules of microbial community invasions. Nature Ecology & Evolution 2019 3:8 3, 1162–1171 (2019).

7. Gilbert, J. A. et al. Current understanding of the human microbiome. Nature Medicine 2018 24:4 24, 392–400 (2018).

8. Turnbaugh, P. J. et al. The Human Microbiome Project. Nature 2007 449:7164 449, 804–810 (2007).

9. Broderick, N. A. & Lemaitre, B. Gut-associated microbes of Drosophila melanogaster. Gut Microbes 3, 307–321 (2012).

10. Obadia, B. et al. Probabilistic Invasion Underlies Natural Gut Microbiome Stability. Current Biology 27, 1999–2006.e8 (2017).

11. Douglas, A. E. The Drosophila model for microbiome research. Lab Anim (NY) 47, 157–164 (2018).

12. Wong, A. C.-N., Chaston, J. M. & Douglas, A. E. The inconstant gut microbiota of Drosophila species revealed by 16S rRNA gene analysis Microbe-microbe and microbe-host interactions. ISME J 7, 1922–1932 (2013).

13. Lesperance, D. N. & Broderick, N. A. Microbiomes as modulators of Drosophila melanogaster homeostasis and disease. Curr Opin Insect Sci 39, 84–90 (2020).

14. Vodovar, N. et al. Drosophila host defense after oral infection by an entomopathogenic Pseudomonas species. (2005).

15. Basset, A., Tzou, P., Lemaitre, B. & Boccard, F. A single gene that promotes interaction of a phytopathogenic bacterium with its insect vector, Drosophila melanogaster. EMBO Rep 4, 205 (2003).

16. Buchon, N., Broderick, N. A., Chakrabarti, S. & Lemaitre, B. Invasive and indigenous microbiota impact intestinal stem cell activity through multiple pathways in Drosophila. Genes Dev 23, 2333 (2009).

17. Chakrabarti, S., Liehl, P., Buchon, N. & Lemaitre, B. Infection-Induced Host Translational Blockage Inhibits Immune Responses and Epithelial Renewal in the Drosophila Gut. Cell Host Microbe 12, 60–70 (2012).

18. Buchon, N., Broderick, N. A., Poidevin, M., Pradervand, S. & Lemaitre, B. Drosophila Intestinal Response to Bacterial Infection: Activation of Host Defense and Stem Cell Proliferation. Cell Host Microbe 5, 200–211 (2009).

19. Gould, A. L. et al. Microbiome interactions shape host fitness. Proc Natl Acad Sci U S A 115, E11951–E11960 (2018).

20. Derke, R. M. et al. The Cu(II) reductase rcla protects escherichia coli against the combination of hypochlorous acid and intracellular copper. mBio 11, 1–17 (2020).

21. Trinder, M., Daisley, B. A., Dube, J. S. & Reid, G. Drosophila melanogaster as a high-throughput model for host-microbiota interactions. Front Microbiol 8, 751 (2017).

22. Chee, W. J. Y., Chew, S. Y. & Than, L. T. L. Vaginal microbiota and the potential of Lactobacillus derivatives in maintaining vaginal health. Microbial Cell Factories 2020 19:1 19, 1–24 (2020).

23. Chen, X., Lu, Y., Chen, T. & Li, R. The Female Vaginal Microbiome in Health and Bacterial Vaginosis. Front Cell Infect Microbiol 11, 271 (2021).

24. O’Hanlon, D. E., Moench, T. R. & Cone, R. A. Vaginal pH and Microbicidal Lactic Acid When Lactobacilli Dominate the Microbiota. PLoS One 8, e80074 (2013).

25. Er, S., istanbullu Tosun, A., Arik, G. & Kivanç, M. Anticandidal activities of lactic acid bacteria isolated from the vagina. Turk J Med Sci 49, 375 (2019).

26. Broderick, N. A., Buchon, N. & Lemaitre, B. Microbiota-induced changes in Drosophila melanogaster host gene expression and gut morphology. mBio 5, 1117–1131 (2014).

27. Blum, J. E., Fischer, C. N., Miles, J. & Handelsman, J. Frequent replenishment sustains the beneficial microbiome of Drosophila melanogaster. mBio 4, 860–873 (2013).

28. Consuegra, J. et al. Metabolic Cooperation among Commensal Bacteria Supports Drosophila Juvenile Growth under Nutritional Stress. iScience 23, 101232 (2020).

29. Henriques, S. F. et al. Metabolic cross-feeding in imbalanced diets allows gut microbes to improve reproduction and alter host behaviour. Nature Communications 2020 11:1 11, 1–15 (2020).

30. latsenko, I., Boquete, J.-P. & Lemaitre, B. Microbiota-Derived Lactate Activates Production of Reactive Oxygen Species by the Intestinal NADPH Oxidase Nox and Shortens Drosophila Lifespan. Immunity 49, 929–942.e5 (2018).

31. Shin, R., Suzuki, M. & Morishita, Y. Influence of intestinal anaerobes and organic acids on the growth of enterohaemorrhagic Escherichia coli O157:H7.

32. Hirshfield, I. N., Terzulli, S. & O’Byrne, C. Weak organic acids: a panoply of effects on bacteria. Sci Prog 86, 245–269 (2003).

33. Lund, P. A. et al. Understanding How Microorganisms Respond to Acid pH Is Central to Their Control and Successful Exploitation. Front Microbiol 11, 2233 (2020).

34. Jugder, B. E., Kamareddine, L. & Watnick, P. I. Microbiota-derived acetate activates intestinal innate immunity via the Tip60 histone acetyltransferase complex. Immunity 54, 1683–1697.e3 (2021).

35. Buchon, N., Broderick, N. A., Poidevin, M., Pradervand, S. & Lemaitre, B. Drosophila Intestinal Response to Bacterial Infection: Activation of Host Defense and Stem Cell Proliferation. Cell Host Microbe 5, 200–211 (2009).

36. Witkin, S. S. et al. Influence of vaginal bacteria and D-and L-lactic acid isomers on vaginal extracellular matrix metalloproteinase inducer: Implications for protection against upper genital tract infections. mBio 4, (2013).

37. Lourenço, A., Pedro, N. A., Salazar, S. B. & Mira, N. P. Effect of acetic acid and lactic acid at low pH in growth and azole resistance of Candida albicans and Candida glabrata. Front Microbiol 10, 3265 (2019).

38. Lesperance, D. N. A. & Broderick, N. A. Gut Bacteria Mediate Nutrient Availability in Drosophila Diets. Appl Environ Microbiol 87, 1–14 (2020).

39. Lesperance, D. N. A. & Broderick, N. A. Meta-analysis of diets used in drosophila microbiome research and introduction of the drosophila dietary composition calculator (DDCC). G3: Genes, Genomes, Genetics 10, 2206–2211 (2020).

40. Adler, P. et al. The key to acetate: Metabolic fluxes of acetic acid bacteria under cocoa pulp fermentation-simulating conditions. Appl Environ Microbiol 80, 4702–4716 (2014).

41. Sommer, A. J. & Newell, P. D. Metabolic basis for mutualism between gut bacteria and its impact on the drosophila melanogaster host. Appl Environ Microbiol 85, (2019).

42. Deshpande, S. A. et al. Acidic Food pH Increases Palatability and Consumption and Extends Drosophila Lifespan. J Nutr 145, 2789–2796 (2015).

43. Fischer, C. et al. Metabolite exchange between microbiome members produces compounds that influence drosophila behavior. Elife 6, (2017).

44. Lee, W. J. & Hase, K. Gut microbiota–generated metabolites in animal health and disease. Nature Chemical Biology 2014 10:6 10, 416–424 (2014).

45. Buchon, N., Broderick, N. A. & Lemaitre, B. Gut homeostasis in a microbial world: Insights from Drosophila melanogaster. Nature Reviews Microbiology Preprint at https://doi.org/10.1038/nrmicro3074 (2013).

46. Lesperance, D. N. A. & Broderick, N. A. Meta-analysis of Diets Used in Drosophila Microbiome Research and Introduction of the Drosophila Dietary Composition Calculator (DDCC). G3: Genes/Genomes/Genetics 10, 2207 (2020).

47. Judd, A. M. et al. Bacterial methionine metabolism genes influence Drosophila melanogaster starvation resistance. Appl Environ Microbiol 84, (2018).

