## Supplemental Figures for "Microbiome derived acidity protects against microbial invasion in *Drosophila*"

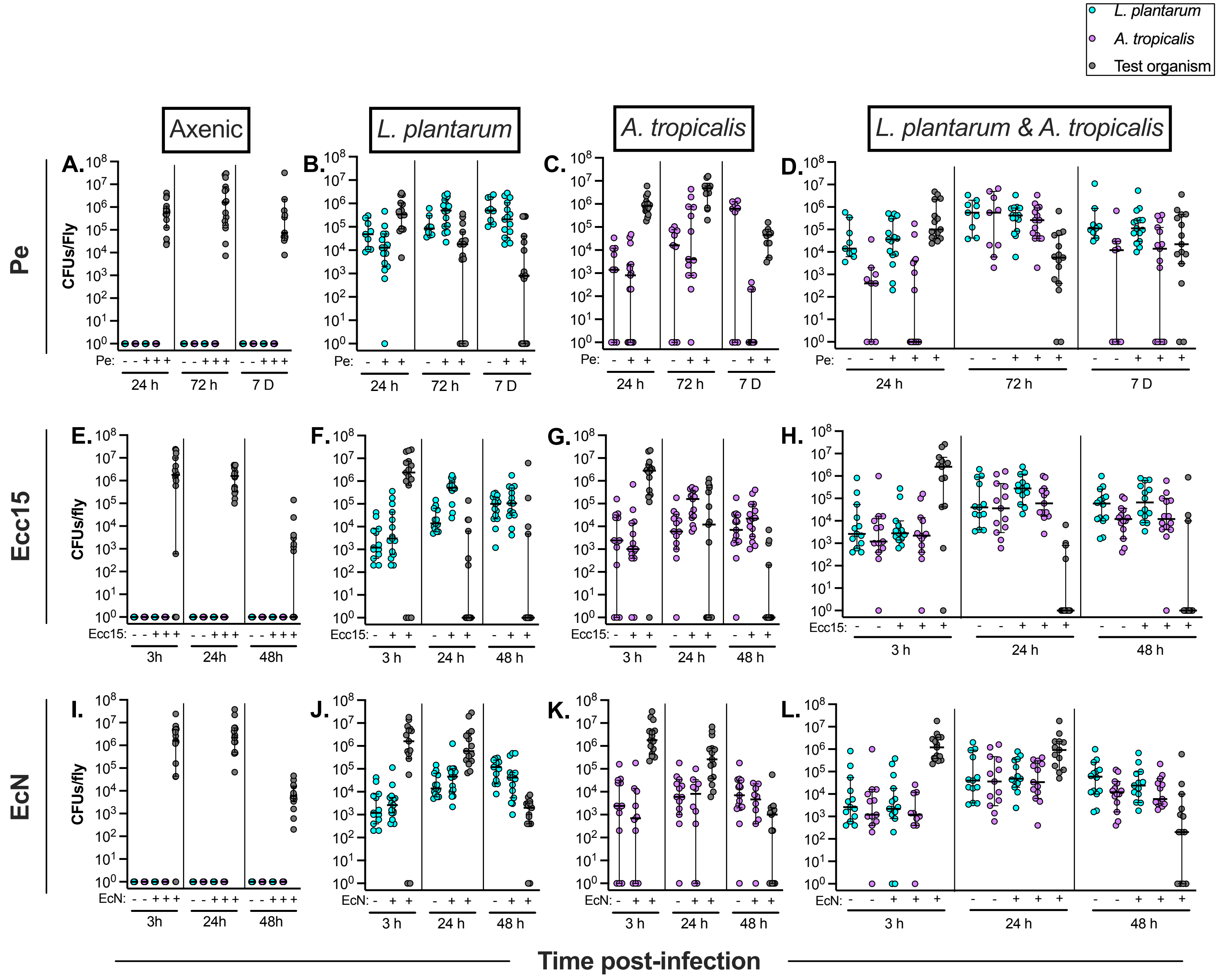


**Figure S1. Microbial Load Analyses of Microbiome Members and Invasive Organisms During Infection.** Number of colony forming units (CFUs) of bacteria per fly infected with Pe (**A-D**), Ecc15 (**E-H**), and EcN (**I-L**). Each invasive organism was screened in female axenic flies and flies colonized with Lp, At, and Lp/At. Flies were sacrificed to determine Pe load at 24 hours, 72 hours, and 7 days post feeding infection (feeding occurred from t=0 to t=24 hours). Ecc15 and EcN bacterial load was determined at 3 hours, 24 hours, and 48 hours post feeding infection. Microbiome load was also assessed at each time point. Each point represents counts from an individual fly; lines and error bars represent the median and 95% confidence intervals; limit of detection is $2x10^2$ CFUs/fly. n=9 flies per control & 10-15 flies (depending on availability of living flies) per infected treatment per time point over 3 biological replicates.

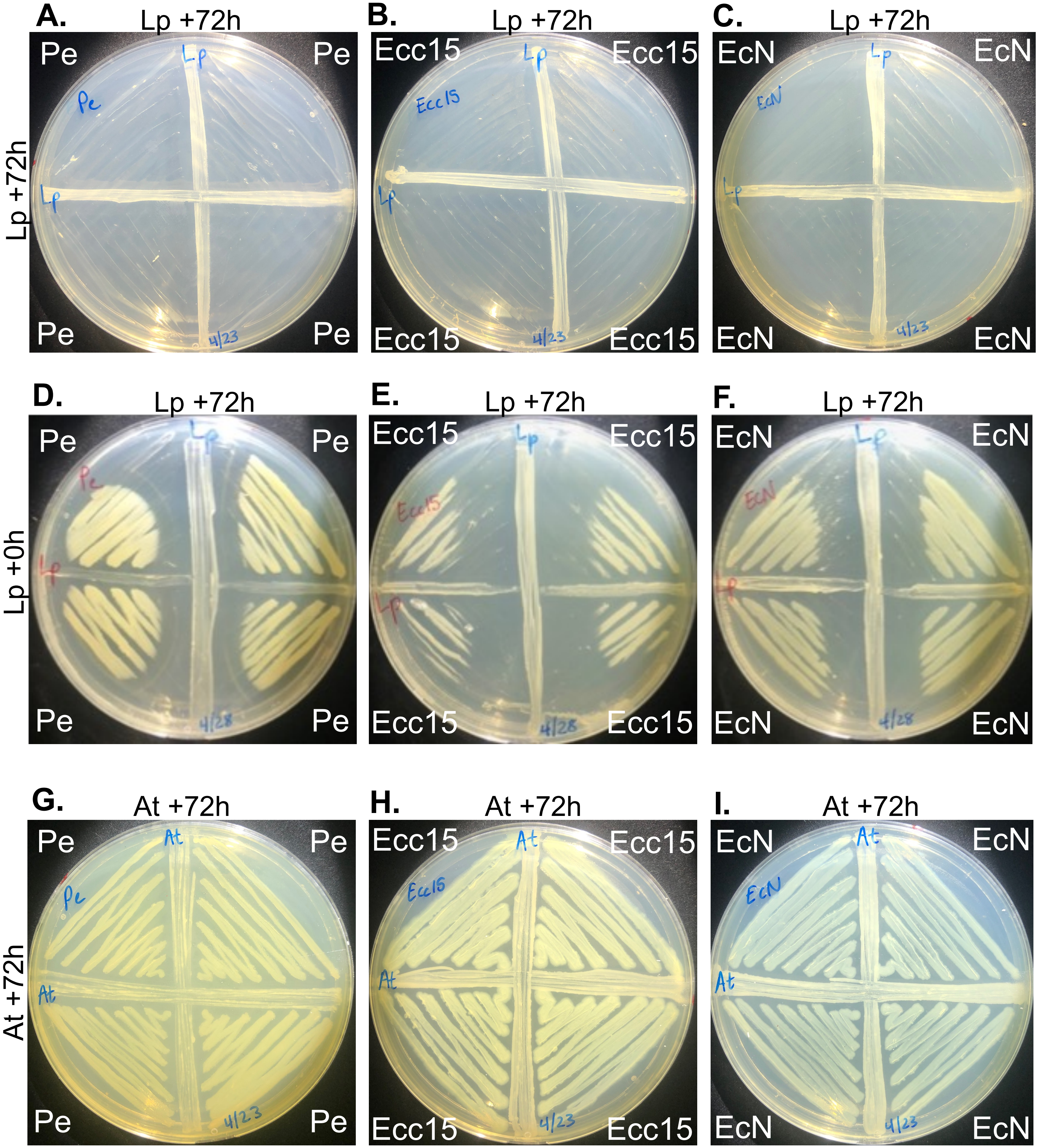


**Figure S2. Lp inhibits organisms in a spatiotemporal manner. A-C)** Multi-organism interactions assays in which Lp is grown on the vertical and on the horizontal for 72 hours, followed by 24 hours of growth of Pe, Ecc15, or EcN in each quadrant. **D-F)** Lp is grown on the vertical for 72 hours. Lp is added to the horizontal at the same time as Pe, Ecc15, and EcN. **G-H)** *A. tropicalis* is streaked on the vertical and horizontal and grown for 72h prior to Pe, Ecc15, and EcN.

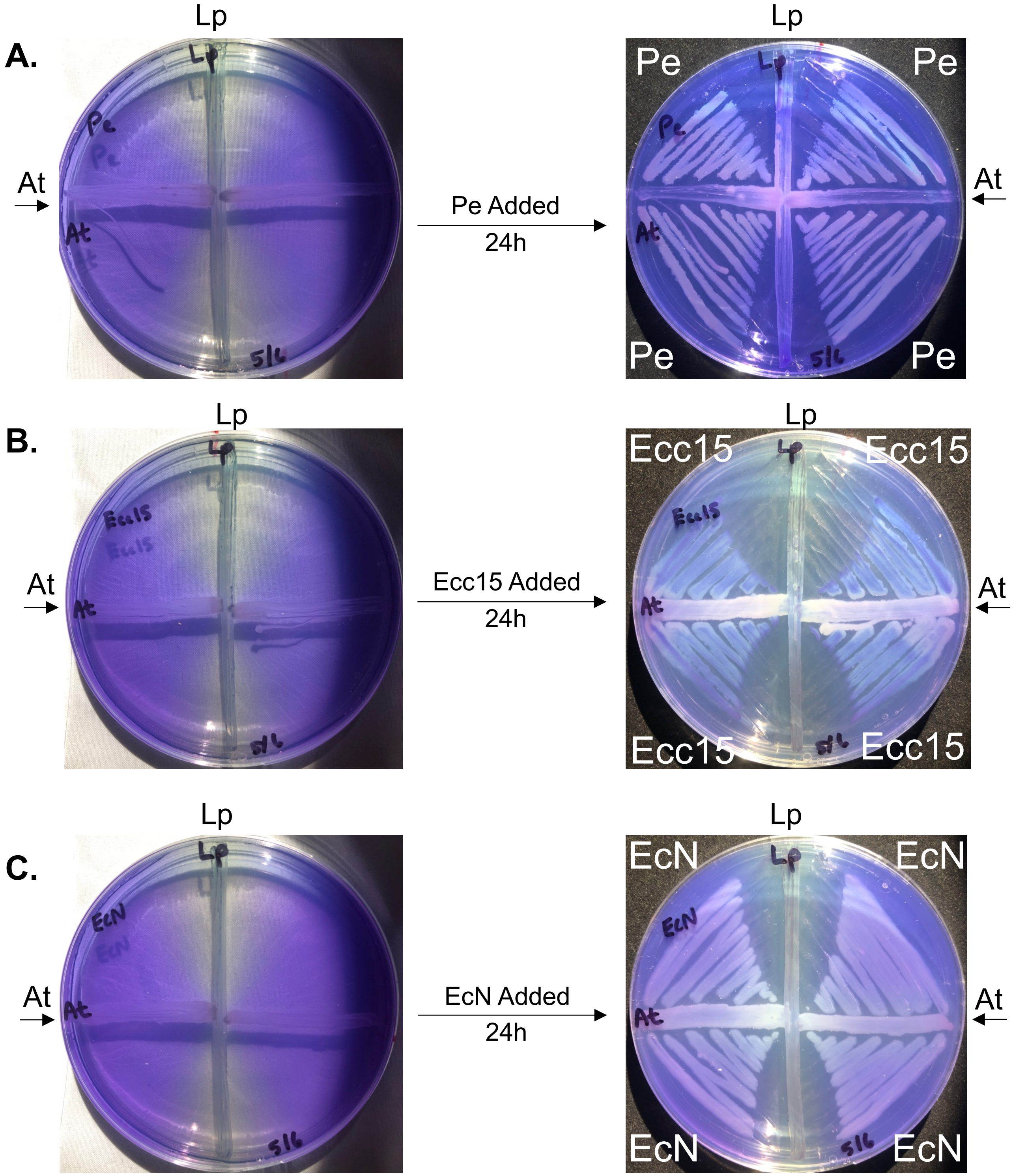


**Figure S3.** Multi-organism interaction assays with the pH indicator bromophenol blue. Lp and At are grown in intersecting lines for three days, after which a yellow acidic zone appears around Lp, but it tapers off near the At interaction point. Invasive organisms Pe **(A)**, Ecc15 **(B)**, and EcN **(C)** are added to the adjacent quadrants and grown for an additional day. Zones of inhibition are then compared to acidic zones.

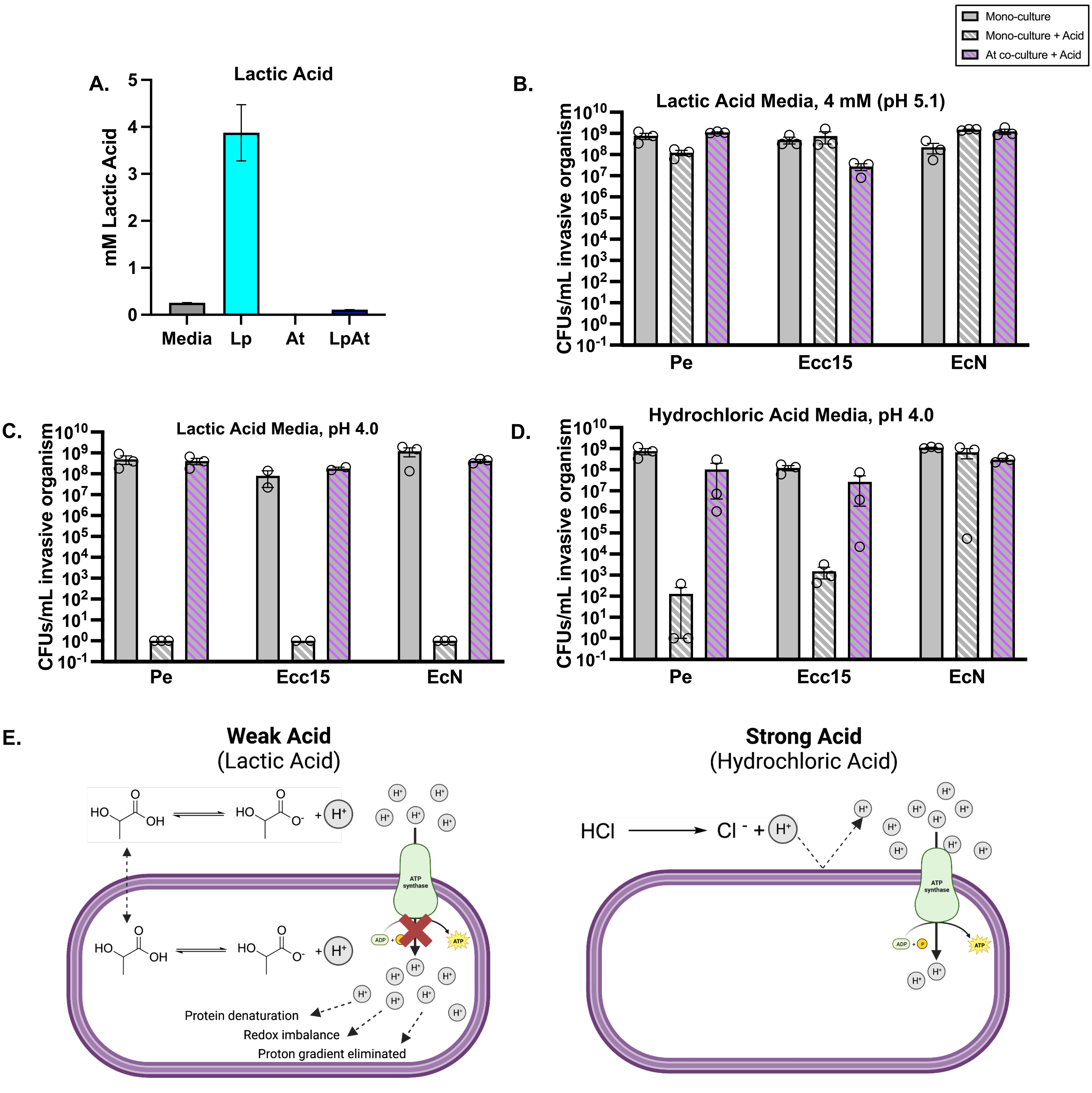


**Figure S4. Microbial inhibition by acid is bifurcated by acid strength and pH. A)** Lactic acid concentrations of overnight cultures of Lp and At and a co-culture of Lp/At. Bars represent mean concentrations of two biological replicates ±SEM. **B)** Microbial concentration of invasive organisms in media supplemented with 4 mM lactic acid (pH ~5.1) with or without 24 hours of prior growth with At. **C-D)** Microbial concentration of invasive organisms in media adjusted to pH 4.0 with lactic acid and hydrochloric acid, respectively, with or without 24 hours of prior growth with At. Each bar represents mean CFUs/mL ±SEM of 3 biological replicates. **E)** Proposed mechanism for differences in antimicrobial activities of weak organic acids and strong acids.

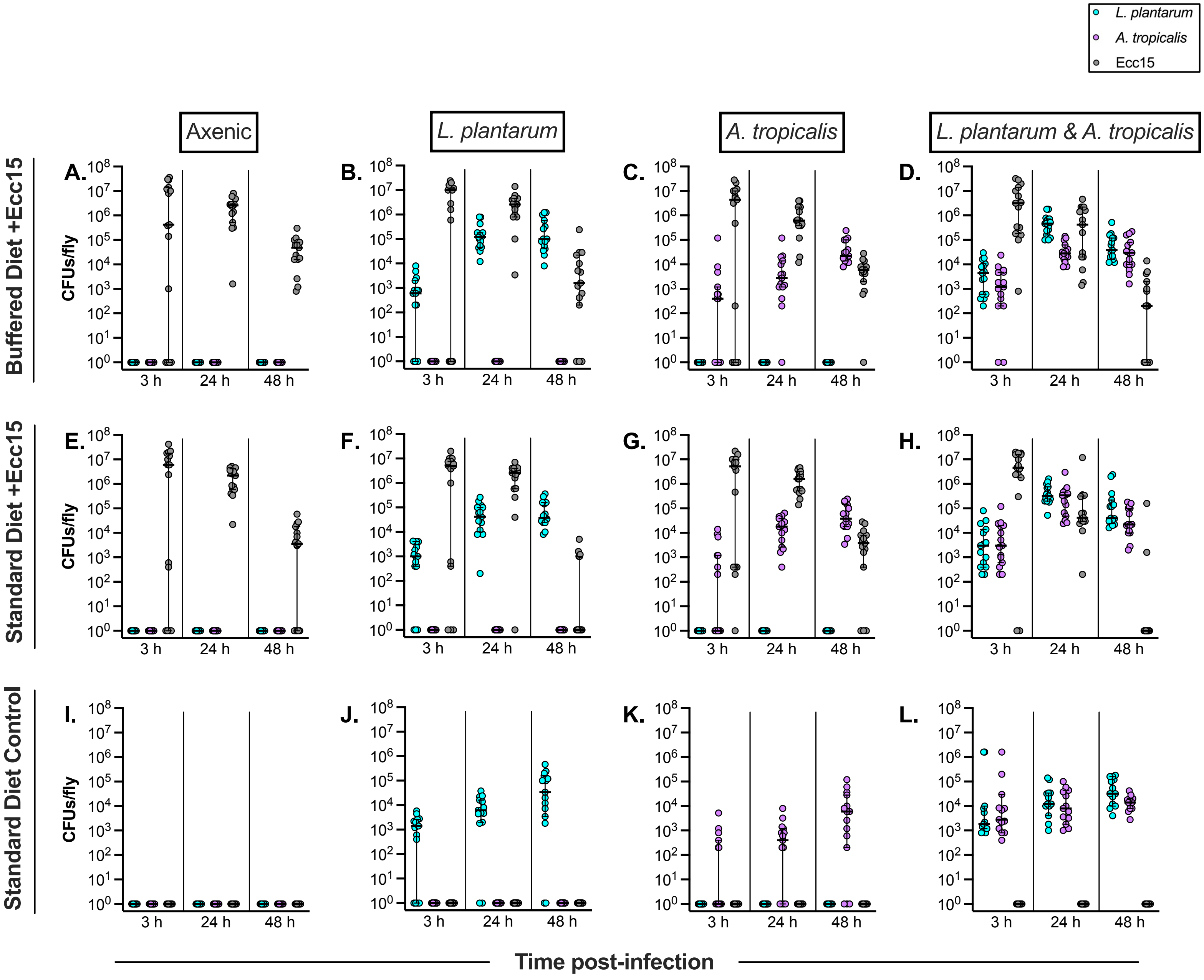


**Figure S5. Microbial Load Analyses of Microbiome Members and Ecc15 During Infections on Buffered and Standard Diets.** Number of colony forming units (CFUs) of Ecc15 and microbiome members (Lp and/or At) per fly infected with Ecc15 on buffered fly diet (**A-D**) and standard fly diet (**E-H**). Uninfected controls were conducted on standard diet (**I-L**). Ecc15 was screened in female axenic flies and flies colonized with Lp, At, and Lp/At. Ecc15 and microbiome bacterial load was determined at 3 hours, 24 hours, and 48 hours post feeding infection. Each point represents counts from an individual fly; lines and error bars represent the median and 95% confidence intervals; limit of detection is $2x10^2$ CFUs/fly. n=10-15 flies per treatment per time point over 3 biological replicates.

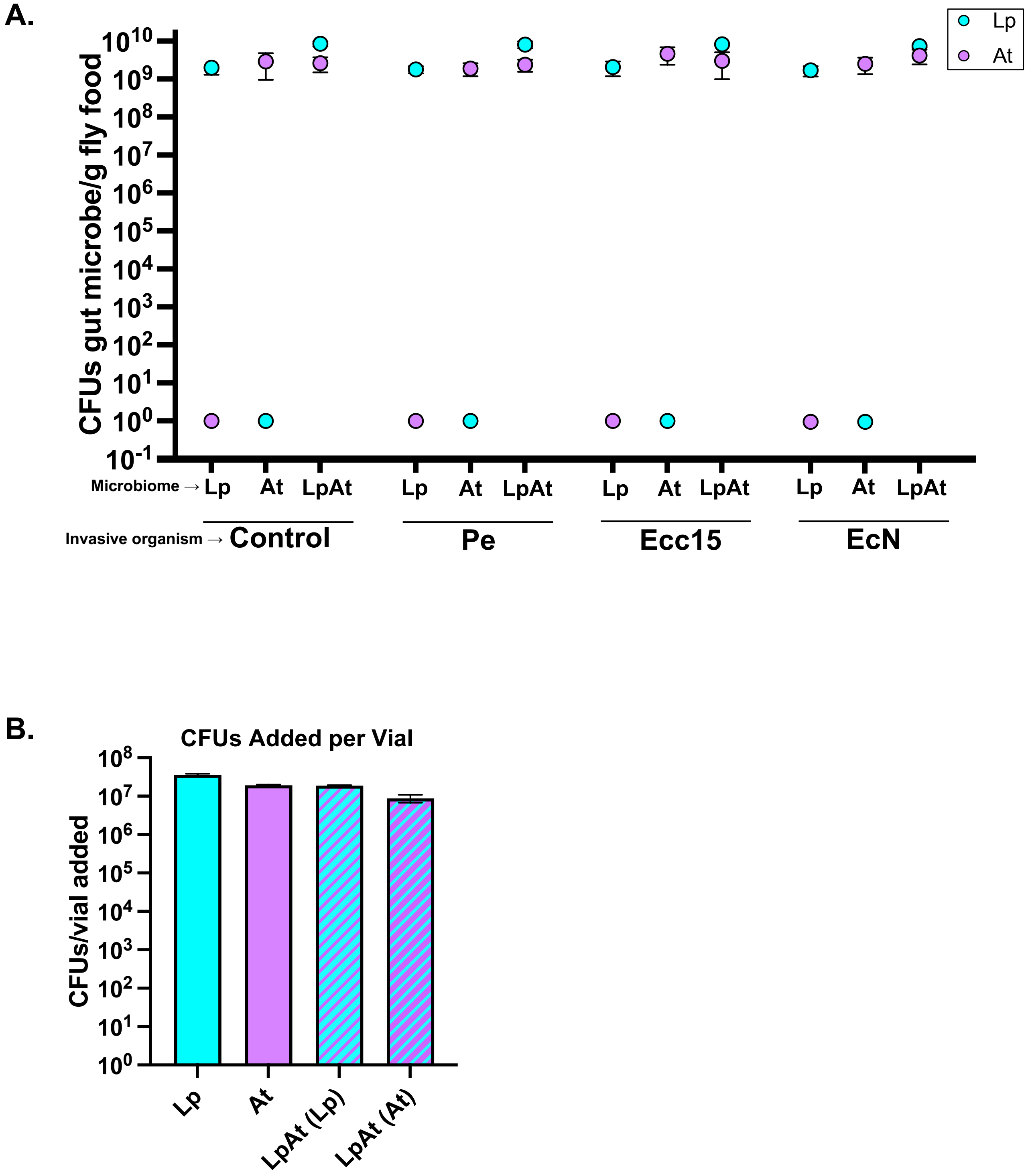


**Figure S6. Levels of Gut Microbe Growth on Fly Food. A)** Bacterial load of Lp and/or At present on fly food after three days of growth followed by an additional day of growth after the addition of invasive organisms (Pe, Ecc15, or EcN). Each point represents the mean CFUs/g fly food ±SEM for three biological replicates. **B)** Number of colony-forming units of each gut microbe in 150 µl of OD 0.5 suspension (the volume added to each vial in **Figure 5C**). Bars represent the mean of three biological replicates ±SEM.
